# Data-driven precision nutrition improves clinical outcomes and risk scores for IBS, depression, anxiety, and T2D

**DOI:** 10.1101/2021.04.24.441290

**Authors:** Janelle Connell, Ryan Toma, Cleo Han-Chen Ho, Nan Shen, Pedro Moura, Tiep Le, Eric Patridge, Grant Antoine, Hilary Keiser, Cristina Julian, Uma Naidoo, Damon Tanton, Guruduth Banavar, Momchilo Vuyisich

## Abstract

Many studies have shown that foods and nutritional ingredients play an important role in healthy human homeostasis, either directly or via the microbiome. We have developed an objective, integrated, and automated approach to personalized food and supplement recommendations that is powered by artificial intelligence and individualized molecular data from the gut microbiome, the human host, and their interactions. The process starts with a clinically validated transcriptomic analysis of a person’s stool (and some cases also blood) sample, from which the following molecular data are computed using clinically validated bioinformatics: 1. Quantitative strain-level taxonomic classification of all microorganisms, 2. Gene expression level for microbial genes, 3. Gene expression level for all human genes. These molecular data are converted into personalized nutritional recommendations (foods and supplements) using algorithms derived from clinical research studies and domain knowledge. We describe an application of our precision nutrition technology platform to human populations with irritable bowel syndrome (IBS), depression, anxiety, and type 2 diabetes (T2D). In these pilot interventional studies, our precision nutrition program achieved significant improvements in clinical outcomes of IBS (39% for severe IBS), depression (31% for severe depression), anxiety (31% for severe anxiety), and the risk score for T2D (>30% reduction relative to the control arm). These data support the integration of data-driven precision nutrition into the standard of care.

## Introduction

There is strong epidemiological and molecular evidence that the human diet contributes significantly to the onset and progression of many chronic diseases and cancers ^1,2,3,4,5,6^. Dietary advice, however, has been controversial, conflicting and polarized. Scientific publications often contradict one another, with one demonstrating health benefits while another suggests potential harm for the same foods or diets ^7,8,9,10^. For example, one epidemiological study reports that daily consumption of fruit juice raises cancer rates significantly ^11^. Other studies report that the consumption of daily fruit or fruit juice can actually prevent cancer ^12,13,14^. Omega 3s have been described as beneficial for mental health, but there are studies that refute such claims ^15,16^. Dairy consumption has also generated contradictory claims, with some evidence supporting an association with a higher incidence of some cancers and chronic diseases, some studies showing no such correlations, while other studies demonstrate significant benefits, and USDA recommends dairy to Americans ^3,17,18,19,20^. Likewise, recent publications provide conflicting data on the effects of saturated fats on cardiovascular and metabolic diseases ^17,18,21,22^. Red meats have been touted as healthy sources of critical nutrients, such as vitamin B12 and dietary protein, and even shown to provide health benefits; yet there is strong epidemiological and mechanistic evidence that they may be responsible for increased rates of inflammatory diseases and cancers ^23,1,24,25^. To complicate the landscape further, current food and supplement guidelines are influenced by the food industry, which promotes the increased consumption of sugars, dairy, and meat ^26,27^.

The large body of conflicting food and supplement information has manifested in the creation of hundreds of diets, each claiming health benefits. This creates confusion and fear in consumers over their health issues or concerns. In addition, traditional dietary assessment methods like food frequency questionnaires depend on the participant to recall their eating habits accurately, and others like food records and 24 hr recalls are long, time consuming, and prone to misreporting due to self-reporting (https://dietassessmentprimer.cancer.gov/profiles/). When it comes to scientific data, the vast majority of food and supplement research does not account for the specific and unique contributions of an individual’s gut microbiome, despite a multitude of evidence showing that the impact is significant ^28,29,30,31,32^. In addition, most dietary guidelines generally focus on food groups (vegetables) rather than single foods (broccoli), assuming that the same food will have the same effect on different people. The USDA-provided nutritional guidelines are the same for everyone and based on population averages (https://www.dietaryguidelines.gov/). These guidelines do not consider personal preferences, food intolerances or food allergies either. Yet the emerging science of the microbiome informs us that each microbiome is like a fingerprint, unique to each one of us ^33^. Now there is strong scientific evidence that genetics plays a minor role, and the gut microbiome plays a major role in the effects of nutrition on human physiology ^34,35^. This was demonstrated in a large twins study of metabolic disease parameters ^31^. This study observed large inter-individual variability in postprandial blood triglyceride, glucose, and insulin responses to identical meals. It is clear that the gut microbiome not only influences how food is digested, but also converts many of the molecular ingredients found in foods into beneficial or harmful secondary metabolites that have profound effects on human physiological functions, such as neurotransmitter production, immune system activation and deactivation (inflammation), immune tolerance, and carbohydrate metabolism ^36,37,38–40^.

Technological advances in genomics and computational biology have made it possible for food and supplement science to adopt a paradigm shift. Each food can now be viewed as a container of molecular ingredients (i.e. micronutrients), rather than a homogenous material that is either good or bad for human health. Most nutritional guidance is offered for generic food groups, nutrients, herbs, spices or even supplements. However nutritional science is moving in a precision-driven way. Hence, the metabolic functions of each person’s unique microbiome need to be identified and quantified, in order to “prescribe’’ specific foods that contain molecular ingredients that will be converted to physiologically beneficial secondary metabolites by that person’s individual gut microbiome. Microbial functions also need to be quantified so each person can avoid foods that contain molecular ingredients that will be converted by their gut microbiome into pro-inflammatory, and therefore harmful secondary metabolites that are associated with disease.

In addition to the approaches that utilize known microbial metabolites and their micronutrient precursors, machine-learnt models from large clinical studies need to be applied to each person’s gut microbiome analysis to guide the best choice of foods and supplements. Such models have already been developed for personalizing foods rich in carbohydrates. These data-driven food choices can minimize the blood glucose levels, especially in the postprandial compartment ^28,30^. Besides the human microbiome, an analysis of human gene expression must be integrated into the systems biology view of the human body to enable the understanding of the network interactions among the dietary molecular ingredients, the microbiome, and human physiology. This understanding is critical because human gene expression, whether modulated by genetic, food and supplement, microbial, or other environmental factors, has a direct connection with the onset and progression of chronic diseases ^41,42,43,44,45^.

In this paper, we describe a data-driven approach to improve clinical outcomes using precision nutrition that are computed by artificial intelligence algorithms using each person’s molecular data and the self-reported phenotype. This approach is completely algorithmic and data-driven, and does not include any opinions, anecdotes, experiences, or pre-determined diets (ketogenic, plant-based, or Mediterranean, for example). The molecular data are obtained from stool or a combination of stool and blood samples using highly accurate and reproducible, clinically validated and licensed transcriptomic tests ^46,47^. These data are used to quantify the activity of microbial metabolic pathways, specific microbial taxa (at the strain level), and human gene expression levels. This information is then converted to personalized foods and supplement recommendations. An overview of this approach is shown in Figure 2 and described in detail in the sections below.

## Materials and methods

### Human subjects, ethical considerations, and study design

The clinical studies described here were approved by a federally-accredited Institutional Review Board (IRB). All samples and metadata were obtained from people at least 18 years old and residing in the USA at the time of participation. All study participants provided informed consent to participating in the studies. The study design was non-conventional; as there was no control arm in three of the studies, due to the limitations of the commercial product. The subjects were blinded to the fact that they were participating in an interventional study with clinical endpoints. All study participants were recruited into a “wellness study” that asked them to complete a wellness survey. These survey questions were clinically validated surveys for depression (PHQ9), anxiety (GAD7), and irritable bowel syndrome (IBS) (Rome IV criteria) ^48,49,50^. Type 2 diabetes (T2D) risk score (0 to 100) was used as the clinical endpoint for T2D ^51^. The T2D trial included two arms, based on the self-reported adherence of each participant, low or high. To evaluate the effect of adhering to the precision recommendations, we compared the difference in T2D risk scores between the “low adherence” cohort (control arm) and the “high adherence” cohort (interventional arm), while controlling for other variables. The two cohorts were matched by (a) initial risk score of +/-10 points (i.e., each participant in one cohort had a participant in the other cohort within 20 points of their risk score at the first time point) and (b) period of adherence of +/- 60 days (i.e., within a total of 120 days), yielding a cohort size of N=1456 for each arm, evaluated over a mean time period of ∼261 +/- ∼132 days and ∼256 +/- ∼133 days for the two arms respectively.

The primary endpoints for the four conditions were:

- IBS: IBS Symptom Severity Score (IBS-SSS), as part of the Rome IV criteria (the Rome Foundation), a clinically validated questionnaire on a scale of 0 through 500, with 300+ indicating severe IBS, 175 to 299 indicating moderate IBS, and 75 to 174 indicating mild IBS ^52^.
- Depression: PHQ9 score, a clinically validated questionnaire of 9 questions that yields a score of 0 through 27, with 15+ indicating moderately severe or severe depression, 10 to 14 indicating moderate depression, and 5 to 9 indicating mild depression ^53,54^.
- Anxiety: GAD7 score, a clinically validated questionnaire of 7 questions that yields a score of 0 through 21, with 15+ indicating severe anxiety, 10 to 14 indicating moderate anxiety, and 5 to 9 indicating mild anxiety ^48^.
- T2D: we used a recently developed T2D risk score, which was built using data from the stool metatranscriptomic analyses of over 50,000 subjects, and validated against a clinical measurement of HbA1c within an independent cohort of over 2200 subjects ^51^.

Each interventional trial had a single arm for participants with depression, anxiety and IBS, and two arms for participants with T2D (high and low adherence). At time point one (T1), participants were asked to complete the wellness questionnaire just prior to receiving their precision food and supplement recommendations. At time point two (T2), each participant was asked to fill out the wellness questionnaire again. There were no additional communications with the participants between the two time points. The clinical studies described here were registered at ClinicalTrials.gov with identifiers NCT04905524 and NCT04905485.

### Metatranscriptomic analyses of stool and transcriptomic analyses of capillary blood samples

The molecular analyses for the studies reported here focus on sequencing messenger RNAs (mRNAs) isolated from human stool and blood samples. Stool samples were collected and analyzed as previously reported ^55^. Briefly, stool samples were collected by the study participants using the Viome commercial kits that included ambient temperature preservation solution and pre-paid return mailers ^47,46^. Stool metatranscriptomic analyses (RNA sequencing, RNAseq) were performed using an automated, clinically-validated laboratory and bioinformatics methods. Results consist of quantitative strain, species, and genus level taxonomic classification of all microorganisms, and quantitative microbial gene and KO (KEGG Ortholog, KEGG = Kyoto Encyclopedia of Genes and Genomes ^56^) expression levels. The matching blood samples were collected and analyzed as previously described ^46^. Briefly, blood samples were collected by the study participants using the Viome commercial kits that included ambient temperature preservation solution and pre-paid return mailers. The kits include lancets and minivettes that enable easy and accurate collection of small volumes of blood (200 uL) from a finger prick. Transcriptomic analyses (RNA sequencing, RNAseq) were performed using an automated, clinically-validated laboratory and bioinformatics methods that require 50 microliters of capillary blood. Test results consist of quantitative human gene expression data.

All microorganisms that live in the intestines obtain their energy by converting chemical substrates into products, using metabolic pathways that consist of enzymes. Substrates are typically the molecular ingredients found in foods, and products are biochemicals usually referred to as secondary metabolites. Metatranscriptomic analysis of the gut microbiome enables the quantification of thousands of microbial pathways using the KEGG database.

### Metabolic pathways and functional scores

We have designed functional scores to quantify certain physiological phenomena; for example, leaky gut, inflammation, gas production, protein fermentation, cellular health, mitochondrial health, etc. These scores are relevant to human physiology and healthy homeostasis. Functional scores are weighted functions (*Score = C*_*1*_*F*_*1*_ *+ C*_*2*_*F*_*2*_ *+ … + C*_*n*_*F*_*n*_, where F is the feature and C is its weight) of specific components from the transcriptomic data. The features or components that make up the functional scores can be taxa, microbial pathways, human pathways, or a combination of other functional scores. Functional scores range from 0 to 100, with 100 representing the highest possible activity. For example, a microbiome-derived functional score such as Butyrate Production Pathways is calculated as a weighted function consisting of the expression levels of many known butyrate-associated KOs (Figure 1) ^57^.

**Figure 1.**
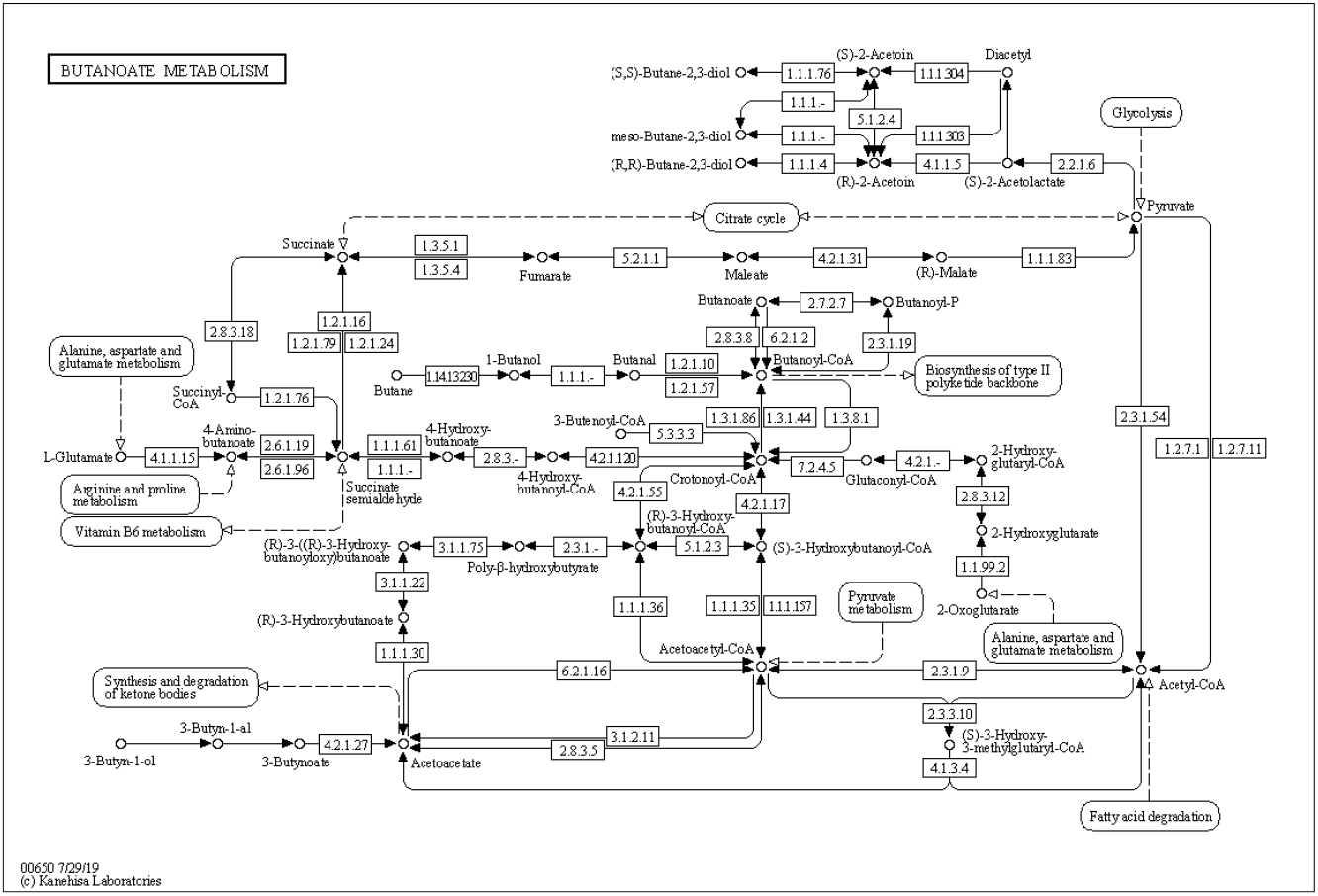
Microbial butanoate (butyrate) production pathways based on KEGG annotations.

The expression level of each KO from the Butyrate Production Pathways score is quantified using the metatranscriptomic stool test and the score is then computed using the weighted formula ^58^. The weights attributed to each KO within a pathway are determined by a combination of domain knowledge (e.g. Figure 1) and statistical analyses obtained from a collection of stool samples ^59,60^. By quantifying each enzymatic member (KO) of the butyrate pathways using microbial gene expression data, estimated pathway activities can be calculated ^61^.

### Precision nutrition program

Viome Precision Nutrition Program (VPNP) includes precision food and personalized supplement recommendations computed by the Viome AI Recommendation Engine. VPNP is designed to maintain the healthy homeostasis at the functional level by increasing beneficial (health-associated) and reducing harmful (disease-associated) biological activities using molecular ingredients from foods and supplements (Figure 2). This approach is built on the concept that on a molecular level, a particular food or supplement is beneficial for one person, but harmful to a different person. Table 1 shows examples of this concept.

**Figure 2.**
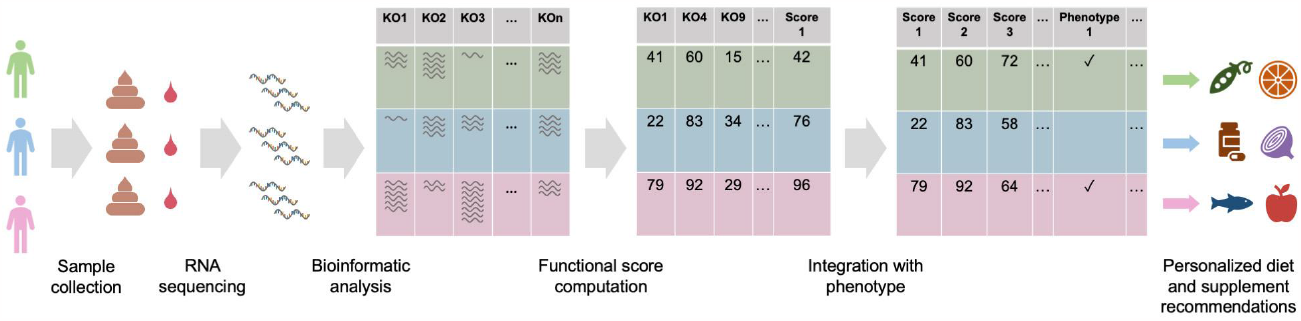
AI-generated precision food and supplement recommendations. Stool and blood samples are collected from participants. The RNA is extracted and sequenced using clinically licensed laboratory and bioinformatics methods. Microbial and human gene expression, including KOs (KEGG orthologs) are quantified from the RNA sequencing data of each participant. Functional scores are subsequently computed as a weighted function of relevant KOs. Finally, personalized nutrition recommendations are computed using all functional scores and phenotypes.

**Table 1:** Examples of specific foods and supplements, and reasons for recommendation to consume or avoid.

|  |  |  |
| --- | --- | --- |
| Food/supplement | Reasons to consume | Reasons to avoid |
| Artichoke | - Low Butyrate Production Pathways functional score, and<br>- Presence of specific microbial strains that convert inulin into butyrate | - High Gas Production functional score, or<br>- Small intestinal bacterial overgrowth (SIBO) symptoms |
| Broccoli | - High Inflammatory Activity functional score, or<br>- Suspected liver and detox support needed | - High Sulfide Gas Production functional score, and<br>- High Trimethylamine (TMA) Production functional score |
| Trout | High Inflammatory Activity functional score | High Uric Acid Production Pathways functional score |
| Nicotinamide Riboside (NR) | - Low Mitochondrial Health functional score, or<br>- Low Mitochondrial Biogenesis Pathways functional score | - High Cellular Senescence functional score, and<br>- High Inflammatory Activity functional score |
| Curcumin | - High Inflammatory Activity functional score, or<br>- High Cellular Stress, or<br>- High Immune System Activation | High Bile Acid Production Pathway functional score |

The Viome AI Recommendation Engine uses a sophisticated set of algorithms to determine the final food and supplement recommendations. The algorithms are developed from domain knowledge (publications on microbial and human physiology, food science, clinical trials, etc.) and extensive clinical studies from which machine-learned models of food and supplement modulation of the microbiome and human physiology were developed ^28^. These algorithms are applied to the functional scores for each person whose stool, or stool and blood samples are analyzed.

The Viome AI Recommendation Engine considers compounds (molecular ingredients) in foods and supplements that can support the molecular functions of health-associated microbial and human genes. These compounds include specific polysaccharides, polyphenols, vitamins, minerals, amino acids, fatty acids, and many phytochemicals. For example, an almond is a source of many phytonutrients and compounds such as kaempferol (flavonoid), naringenin (flavonoid), ferulic acid (phenolic acid), oxalic acid, phytic acid, quercetin, procyanidin B2 and B3, magnesium, phytosterols such as retinol, a-tocopherol, vitamin K, vitamin D, and beta-sitosterol, fatty acids such as oleic acid, linoleic acid, and palmitic acid, and specific amino acids ^62^. The decision to recommend a specific compound and its amount depends on the values of multiple functional scores. After considering all inputs, the recommendation engine classifies foods into one of four categories, based on the molecular composition of each food. The food categories are superfoods, enjoy foods, minimize foods, and avoid foods, which are consumerized names that correspond to the recommended servings per day for each food.

Personalized supplement recommendations follow the same logic, considering all inputs to identify compounds that are beneficial or harmful to an individual’s functional scores and phenotype. Supplements include minerals, vitamins, botanicals or herbs, food extracts, enzymes, phospholipids, amino acids, prebiotics, and probiotics. When considering individual functional scores, supplement ingredients commonly believed to be beneficial may not be recommended. For example, turmeric is a commonly consumed supplement for its anti-inflammatory properties, but has also been shown to increase bile flow ^63,64,65^. While this was an animal study, it can help to inform us that for individuals with a high Bile Acid Metabolism Pathway functional score, turmeric supplementation may be more harmful than beneficial and must be assessed more carefully. A high Bile Acid Metabolism Pathway score suggests that the microbial activity of transforming bile salts into bile acids is high. While such biotransformation is part of a balanced gut microbiota and bile acid homeostasis, excessive intestinal bile acids may promote a pro-inflammatory environment and play a role in the development of gastrointestinal diseases ^66,67^.

The process of categorizing foods and supplements (determining the servings or dose) includes prioritizing scores that need improvement and considering conflicts within the recommendations. An example is shown in Table 2: a low Energy Production Pathway functional score will yield recommendations based on compounds that contribute to the score activity, one of which is alpha-lipoic acid (ALA). Spinach and broccoli are recommended due to the ALA content that is a critical cofactor for mitochondrial energy-production enzymes such as pyruvate dehydrogenase (PDH), alpha-ketoglutarate dehydrogenase (alpha-KGDH), and branched-chain ketoacid dehydrogenase (BCKDC) ^68^. However, when considering additional score results, broccoli and spinach are placed on the avoid food list due to broccoli’s glucosinolate content and spinach’s oxalate content. Instead, tomatoes and peas are recommended as sources of ALA to support the Energy Production Pathway functional score.

**Table 2:** Food Recommendations Case Example

| Table 2: Food Recommendations Case Example |  |  |  |  |
| --- | --- | --- | --- | --- |
| Example Score Results:<br>High Sulfide Gas Production Pathways functional score<br>Low Butyrate Production Pathways functional score<br>Low Oxalate Metabolism Pathways functional score<br>Low Energy Production Pathways functional score |  |  | Self-reported information on wellness questionnaire:<br>Histamine intolerance<br>Pistachio allergy |  |
| Food | Compounds | Food categories (initial) | Final recommendation | Replacement food/supplement |
| Spinach | - Alpha-lipoic acid<br>- Oxalates | - Superfood, based on low Energy Production Pathways functional score<br>- Avoid, due to low Oxalate Metabolism Pathways functional score | Avoid | - Tomato and<br>- Alpha-lipoic acid supplement |
| Broccoli | - Alpha-lipoic acid<br>- Glucosinolates | - Superfood, based on low Energy Production Pathways functional score<br>- Avoid, due to high Sulfide Gas Production Pathways functional score | Avoid | - Peas and<br>- Alpha-lipoic acid supplement |
| Sauerkraut | Probiotics | - Superfood, based on low Butyrate Production Pathways functional score<br>- Minimize, due to self-reported histamine intolerance | Minimize | - Probiotic supplement that includes *Lactobacillus acidophilus*, *Lactobacillus plantarum*, *Lactobacillus rhamnosus* |
| Pistachios | CoQ10 | Superfood for Energy Production Pathways Low<br>Avoid due to self-reported pistachio allergy | Avoid | CoQ10 supplement |
| Grapes | Resveratrol | Superfood for Energy Production Pathways Low | Superfood |  |

There are circumstances where a beneficial compound cannot be obtained from food due to an allergy or other health-related issue, or due to the lack of sufficient amounts in food. Personalized supplements help support those gaps in nutrition. In Table 2, pistachios are recommended due to their CoQ10 content. However, if an individual has an allergy to pistachios or exhibits small intestinal bacterial overgrowth (SIBO) symptoms, pistachios are placed on the avoid or minimize food list. In this situation, CoQ10 can be provided through a supplement as the recommendation engine associates it as beneficial for the same score and/or phenotypic conditions.

## Results

We conducted four interventional studies that tested the efficacy of VPNP to improve clinical or molecular scores for IBS, depression, anxiety, and T2D. Three of the studies included only an interventional arm (IBS-SSS, n=105; PHQ-9, n=410; GAD-7, n=490); the T2D study also included a control arm (1456 low adherence vs. 1456 high adherence).

Figures 3A, 3B, and 3C show the changes in the clinical study endpoints following VPNP. We observe an improvement in the clinical scores for all three conditions (IBS, depression, and anxiety) and all disease activity levels. We note that the p-value for mild depression score improvement is 0.2038, which is above the conventional standard of 0.05. However, the improvement in all other clinical outcomes, conditions and severity levels are highly statistically significant.

**Figure 3:**
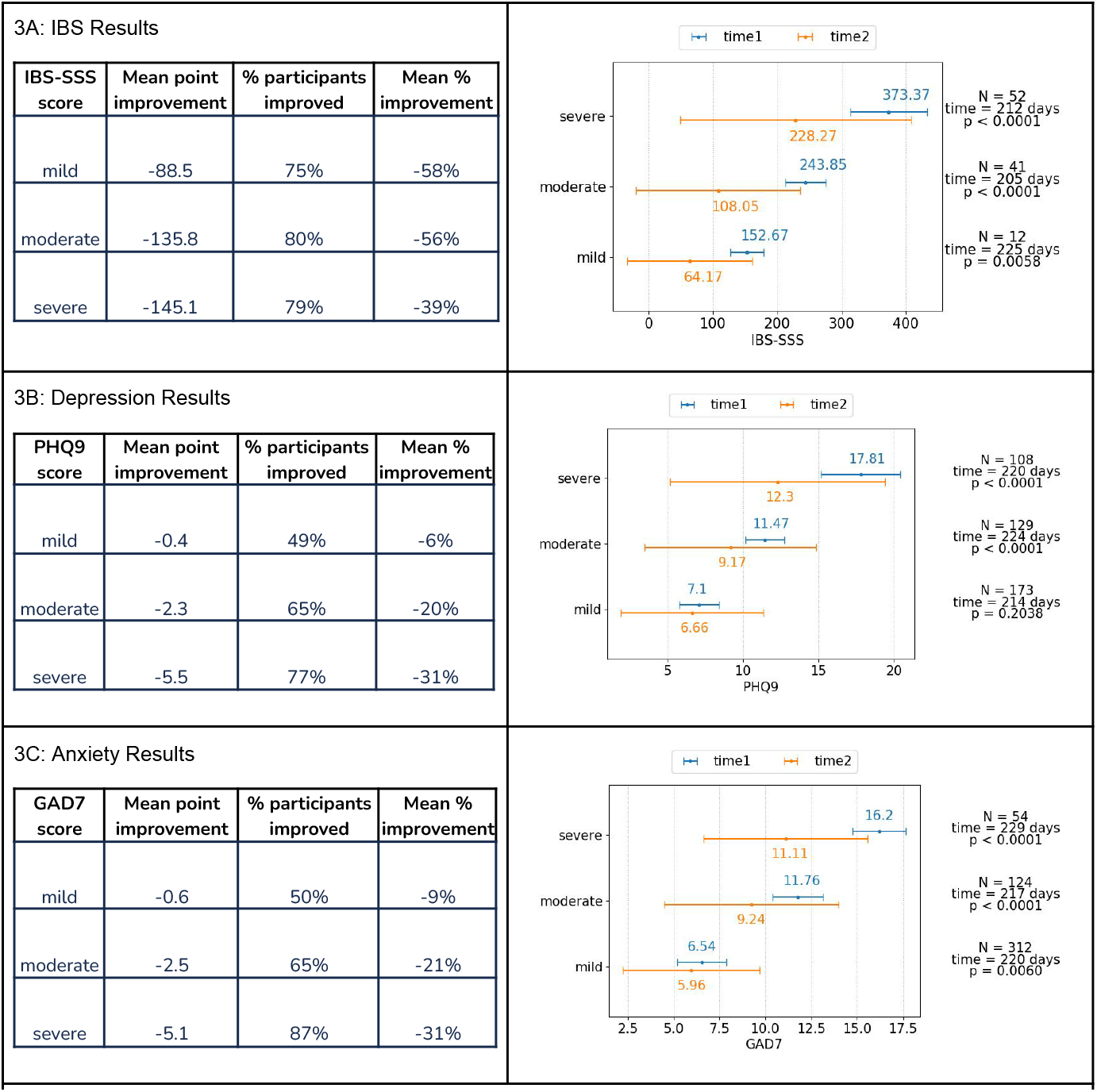
Efficacy of the food and supplement interventions on participants with IBS, depression, or anxiety. A. Results showing the effect of precision food and supplement recommendations on subjects with IBS, measured using the IBS-SSS score. Bars on the right are the mean +/- standard deviation of IBS-SSS score; blue bars represent the baseline (at the study start) and orange bars represent the followup (after the nutritional intervention). B. Results showing the effect of precision food and supplement recommendations on subjects with depression, measured using the PHQ9 score. Bars on the right are the mean +/- standard deviation of PHQ9 score; blue bars represent the baseline (at the study start) and orange bars represent the followup (after the intervention). C. Results showing the effect of precision food and supplement recommendations on subjects with anxiety, measured using the GAD7 score. Bars on the right are the mean +/- standard deviation of GAD7 score; blue bars represent the baseline (at the study start) and orange bars represent the followup (after the intervention).

Figure 4 shows the difference in the change of the T2D risk score (y axis) between the low-adherent participants (blue box plot) and high-adherent participants (green box plot). We observe a statistically significant risk score improvement for people with high adherence (p=1.99e-05), with a mean risk score reduction of 7 points [(−30.25) - (−23.21)], which translates to >30% improvement. This shows that high adherence to the precision nutrition recommendations results in a lower risk of T2D.

**Figure 4.**
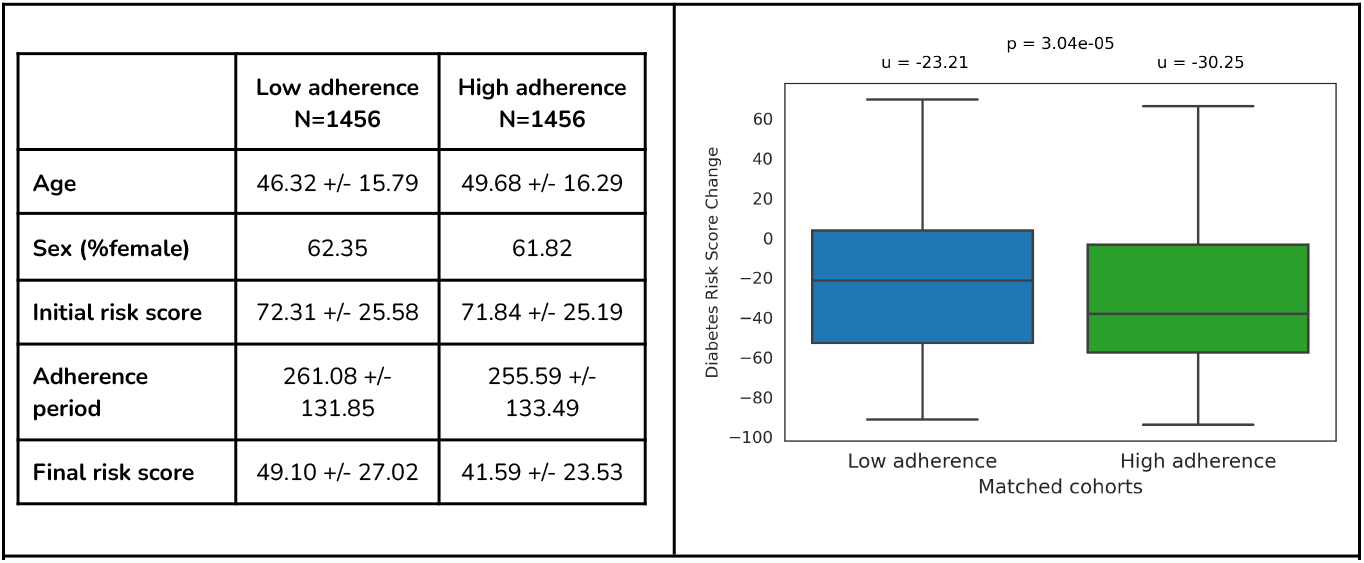
Efficacy of the food and supplement interventions on participants with an elevated T2D risk score. Participants who adhered to precision recommendations (green box) reduced their T2D risk score more than participants who did not (blue box) (p<0.001).

## Discussion

We describe a novel technology platform that overcomes some of the major shortcomings of food and supplement science, and improves clinical and molecular outcomes in several highly prevalent chronic diseases, IBS, depression, anxiety, and type 2 diabetes ^69,70,71^. Traditional food and supplement research has been conducted using various methods. Epidemiological studies have been used to derive food and supplement rules using large populations. These have yielded interesting results that have guided certain national-scale dietary recommendations, such as the current USDA guidelines in the USA (Dietary Guidelines for Americans, 2020-2025). Many clinical and pre-clinical studies have also tested the effects of certain foods and diets on various health outcomes. Additionally, many different supplements, including prebiotics and probiotics, and their combinations have been tested for their ability to improve certain symptoms ^72,73,74,75,76,77^.

Over the last decade, it has become clear that the gut microbiome plays a crucial role in modulating human physiology by regulating its immune system, metabolic functions, hormones, and neurotransmitters ^78^. It is important to note that the microbiome’s influence on human physiology is exerted via its functions, and not merely the taxonomy ^79,80^. For example, the fact that a person’s gut microbiome contains *Faecalibacterium prausnitzii*, which is a well-known butyrate producer, does not mean that it is producing butyrate. However, when it encounters the correct substrate, it will produce this important metabolite ^81,82^. *Escherichia coli* is a bacterial species that comprises very beneficial strains that produce vitamins K and B12, and help human hosts efficiently absorb iron; it also represents highly pathogenic, life-threatening microorganisms, such as the enterohemorrhagic strains ^83,84,85^. Therefore, identifying the species of *Escherichia coli* in a gut microbiome is not very informative, given the vastly different functions that members of this species can perform.

The key to understanding the effects of the gut microbiome on human physiology and health is to quantify the microbial functions using metatranscriptomic, metaproteomic, or metabolomic approaches. We have chosen metatranscriptomics as the best single approach, as it enables quantitative measurements of microbial gene expression that can identify both protein-based and metabolite-based effects on the human physiology, and still provide the highest resolution of taxonomic classification (can distinguish strains) that can guide some aspects of the food and supplement recommendations. Another essential aspect to understanding the relationship between the human physiology, microbiome and diet is to integrate the human molecular data into the systems biology view of the human body. We use a whole blood transcriptome test that can quantify the expression levels of >10,000 human genes from capillary (finger prick) blood.

We also want to emphasize that foods should be viewed as containers of molecular ingredients, instead of objects that we can visually recognize as onions, peppers, etc. The reason for this is that each food contains many different molecular ingredients, as exemplified above, that can exert a multitude of health-related effects on the host, either directly or via the microbiome metabolism ^86^.

Study limitations: Three of the four studies (for IBS, depression, and anxiety) were single arm interventional studies without control arms. We also did not capture the level of adherence to the food and supplement recommendations for these three studies, which could significantly affect the observed results; improved adherence may have provided improved efficacy. We did capture this information for the T2D study, and showed that improved adherence improved the molecular scores (Figure 4). It is possible that the observed reductions in the clinical scores would have been higher if the participants with low adherence were excluded from the study. For the T2D study, we matched the study arms for age, sex, and BMI, but did not include other possible confounders. The diet and supplement recommendations were delivered digitally, and no effort was made to either monitor or improve compliance. Because these are preliminary studies without robust control arms, the observed improvements could be explained by the regression to the mean. All of these limitations will be addressed in future randomized controlled trials.

## Notes

### Competing Interest Statement

Most authors are employees of Viome Life Sciences, Inc., the sponsor of the research.

### Summary of Updates

We expanded the study to include a larger cohort of participants that joined the study since the first version was published. We also added a few authors that contributed to the data and manuscript.

